# Multitasking boosts muscular endurance task performance due to elevated arousal level unattainable by the endurance task alone

**DOI:** 10.64898/2026.03.06.710139

**Authors:** Sofia Nagisa, Ethan Oblak, Shinsuke Shimojo, Kazuhisa Shibata

**Author notes:** Correspondence (K.S.).

## Abstract

Multitasking is generally regarded as detrimental to performance. This deterioration effect is typically explained by the interference among tasks due to the limited capacity of information-processing resources, which in turn reduces the performance in each task. Contrary to this general view, we report evidence for a facilitation effect of multitasking on performance. This facilitation effect was observed in multitasking on a handgrip muscular endurance task and cognitive task, which are known to have little interference with each other. Specifically, we found that performance in the endurance task was facilitated with the difficulty of the concurrent cognitive task. This facilitation effect was mediated by additional pupil dilation due to the cognitive task. Increased effort with the difficulty of the cognitive task cannot explain the facilitated performance in the irrelevant endurance task. Instead, they suggest that the cognitive task elevated overall arousal to a level unattainable by the endurance task alone, which in turn facilitated performance in the irrelevant endurance task. To further test this arousal account, we manipulated participants’ motivation to the cognitive task by reward without changing its difficulty and found the same pattern of results. Thus, it is not effort or motivation specific to the cognitive task but rather overall arousal level that underlies the facilitation effect. These results unveiled a previously overlooked mechanism: a multitasking-induced arousal boost. Our findings suggest that multitasking can facilitate performance when the net effect of adding a concurrent task is governed less by the capacity limitation and more by the elevation of overall arousal.

## Introduction

One of the main goals of cognitive science is to identify the factors that influence our performance and establish ways to improve it. Multitasking, here defined as concurrently executing two tasks, has become increasingly important as a subject of research in this regard, since most people in the current world engage in multitasking. In general, multitasking impairs performance, for example when using their smartphones while walking^1^. But, is it possible that multitasking could facilitate performance?

In many tasks, including perceptual recognition^2^, cognitive control^3^, memory^2,4–9^, motor tasks^3,5,8,10,11^, and more complex tasks like driving^12^, performance suffers from multitasking. This is often explained by interference due to the limited capacity of information-processing resources among multiple tasks, which reduces the performance in each task^13–16^. This prior research has established the prevailing view that multitasking typically impairs performance.

However, the same line of research also proposed that a global brain state such as arousal^17,18^ can increase the capacity^13,15^. It is well known that arousal is elevated by the execution of a task^13,19–24^. Collectively, these previous findings raise a new possibility: multitasking facilitates performance through the elevation of overall arousal due to the execution of multiple tasks.

Nevertheless, a facilitation effect of multitasking has been hardly reported. One potential reason is that a deterioration effect masked a potential facilitation effect in previous multitasking studies. However, in some pairs of tasks involving motor execution and simple cognitive processing, performance is relatively unaffected by multitasking^24,25^. Thus, a motor-cognitive task pairing provides a useful testbed for examining a facilitation effect by multitasking. In such a pair, the net effect of adding a concurrent task may be governed less by the interference between tasks and more by the increased capacity due to arousal elevation.

Based on this idea, the present study tested the possibility of a facilitation effect on performance by multitasking. Specifically, we conducted a series of behavioral experiments to systematically assess how performance in a handgrip muscular endurance task was affected by a concurrent cognitive task. Results showed that the endurance force performance increased with the difficulty and amount of reward assigned to the cognitive task. This increased force was mediated by additional pupil dilation due to the cognitive task, consistent with the crucial role of arousal^19–21^. On the other hand, increased effort and motivation to the cognitive task cannot explain these effects because they should be specific to the cognitive task while irrelevant to the endurance task^26–36^. These results suggest a mechanism in which the concurrent cognitive task elevated overall arousal to a level unattainable by engaging in the endurance task alone, thereby facilitating performance in the endurance task.

Our findings present evidence of a facilitation effect that multitasking has on performance, calling for a reconsideration of the prevailing view claiming that multitasking is inherently detrimental to performance.

## Results

### Performance in a handgrip endurance task was facilitated by a demanding concurrent cognitive task

In Experiment 1 (N = 24), we tested whether the concurrent execution of two tasks could lead to facilitation of performance. Specifically, we examined whether performance on the focal task is facilitated with the difficulty of the concurrent task. Increasing the difficulty of the concurrent task likely recruits greater attention and effort toward that task^26–30,35^. If the increased concurrent-task difficulty elevates a nonselective global brain state (e.g., arousal^17,18^), we should observe a facilitation effect on focal-task performance.

We deliberately designed our experiments in a way that allows us to test facilitation effects while minimizing deterioration effects due to interference. Previous studies mostly focused on deterioration effects due to interference, which arises from the limited capacity of information-processing resources for multiple tasks^13–16^. Such interference-induced deterioration effects would likely cancel out any potential facilitation effects.

Thus, we selected a set of tasks that were previously demonstrated to have little interference, namely a simple motor task and cognitive task^25,37^. As a motor task, we employed a muscular endurance task in which participants were asked to grip a force sensor using one arm with the highest force level they could sustain for 12 seconds (Figure 1A, right; see Methods for details). In a concurrent cognitive task, participants were presented with a sequence of object images (Figure 1A, left).

**Figure 1.**
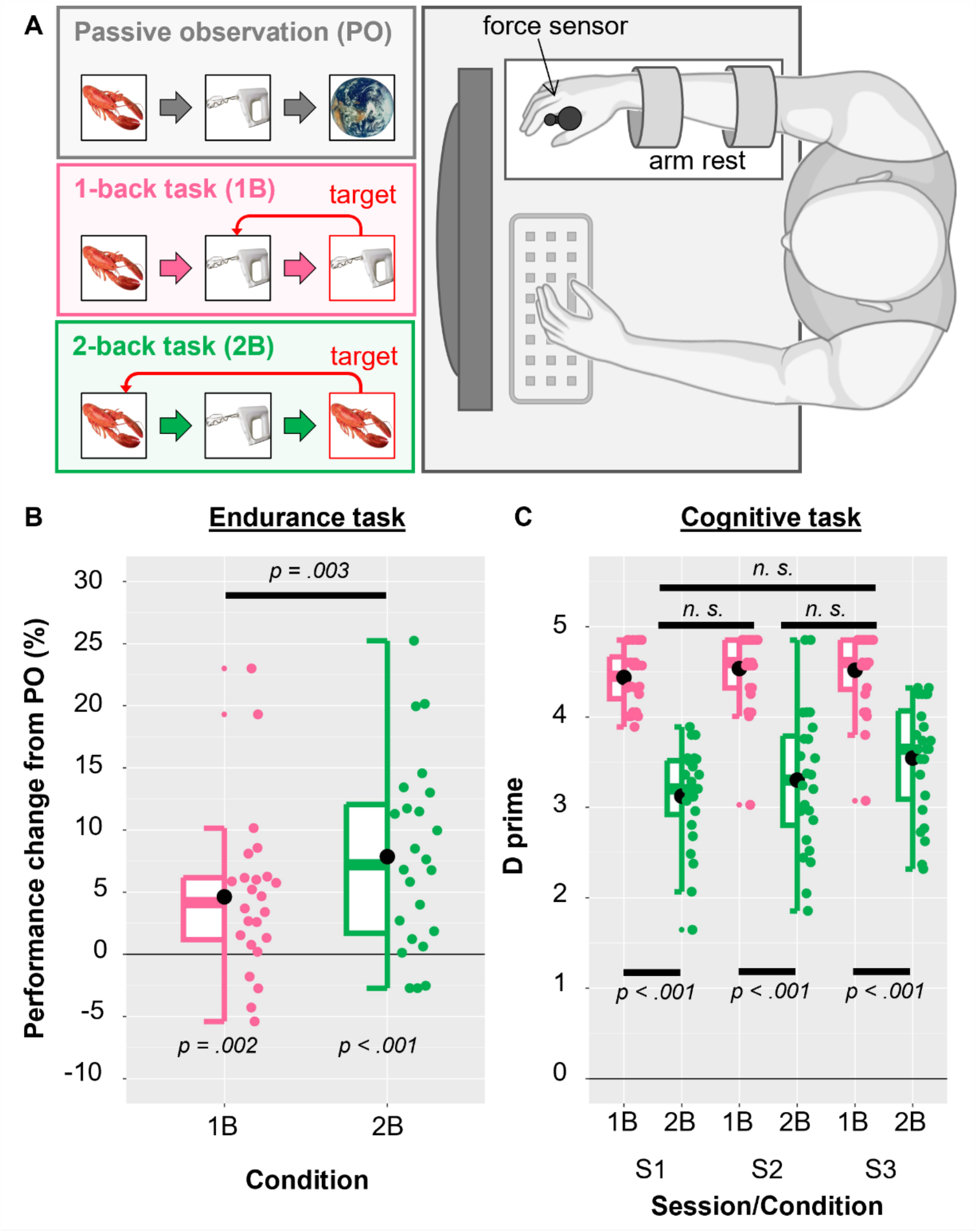
Procedures and results of Experiment 1. (A) Multitask setting in Experiment 1. Participants performed a handgrip muscular endurance task (right) while simultaneously engaging in a cognitive task under one of the three conditions (left): passively observing images (PO), performing 1-back task (1B), and performing 2-back task (2B) (see Methods for details). (B) Percent changes in endurance performance in the 1B and 2B conditions, relative to the PO condition as a baseline. Average performance changes was significantly larger than zero in the 1B and 2B conditions. In addition, performance change was significantly higher in the 2B condition than in the 1B condition. (C) Performance in the cognitive task. Detection sensitivity (d’) did not significantly differ among multitasking sessions (S1-S2) and single task session (S3). Box plots show the medians, lower and upper quartiles, and minimum and maximum adjacent values without outliers across participants. Black dots represent the mean values across participants, and colored dots represent individual values.

To manipulate participants’ engagement in the cognitive task, we introduced three conditions commonly used for such manipulation^38^: passive observation (PO), 1-back task (1B), and 2-back task (2B), with difficulty increasing in this order (see Methods for details). On each trial, participants performed the endurance task while engaging in one of the three conditions of the cognitive task. In the first session of multitasking (30 trials), they used one arm for the endurance task and the other arm to make a button press for the cognitive task. In the second session (30 trials), the arms were switched. Performance in the endurance task was defined as the mean force exerted during the task (see Methods for details). Performance in the 1B and 2B conditions of the cognitive task was defined as sensitivity (d’) of target detection.

Results of the endurance task are shown in Figure 1B as percent change of exerted force in the 1B and 2B conditions relative to the PO condition as a baseline. We used this performance change metric to assess the effect of cognitive task difficulty on endurance performance because of substantial inter-individual variability in exerted force. To compare the performance changes between the two conditions of the cognitive task, we conducted a two-way repeated-measures ANOVA with Condition (1B vs. 2B conditions) and Session (first multitasking vs. second multitasking sessions) as factors. We found a significant effect of Condition (*F*_2,46_ = 11.097, *p* = 0.003) on performance changes, but no significant effect of Session (*F*_1,23_ = 0.014, *p* = 0.907) or interaction between the factors (*F*_2,46_ = 0.337, *p* = 0.567). In addition, results of subsequent tests (one-sample t-test with Bonferroni correction for multiple comparisons) indicated significant performance changes for the 1B condition (*t*_23_ = 3.501, corrected *p* = 0.004) and 2B condition (*t*_23_ = 5.090, corrected *p* < 0.001). These results suggest that the endurance performance was facilitated with the level of engagement in the concurrent cognitive task.

The lack of significant interaction between Condition and Session was commonly observed in all other experiments in this study. Thus, we averaged data in the endurance task across the first and second multitasking sessions throughout this study.

One may ask whether there was an analogous facilitation effect on cognitive task performance due to the endurance task. To test this possibility, after the multitasking sessions we introduced a single task session in which participants performed only the cognitive task. If the possibility were indeed the case, we should expect significant difference in performance of the cognitive task between the multitasking and single task sessions. We conducted a two-way repeated-measures ANOVA (Figure 1C) with Session (first multitasking, second multitasking, vs. single task sessions) and Condition (1B vs. 2B conditions) as factors. As expected, a significant effect of Condition (*F*_1,23_ = 113.345, *p* < 0.001) was observed, confirming that the difficulty in the 2B condition was indeed higher than that in the 1B condition. We also found a significant effect of Session (*F*_2,46_ = 3.553, *p* = 0.037), but not a significant interaction between the factors (*F*_2,46_ = 2.403, *p* = 0.102). However, results of the subsequent tests (paired t-test with Bonferroni correction for multiple comparisons) showed no significant differences in performance of the 2B condition for any combinations among the three sessions (*t*_23_ values < 2.463, *p* values > 0.065). Thus, no or little facilitation effect of multitasking occurred in the cognitive task.

The results of Experiment 1 demonstrated that multitasking can indeed facilitate performance. In particular, engaging in the more demanding cognitive task led to facilitated performance in the muscular endurance task. Increased effort with the difficulty of the cognitive task cannot explain the facilitated performance in the irrelevant endurance task. Rather, they suggest that the cognitive task elevated overall arousal to a level unattainable by the endurance task alone, which in turn facilitated performance in the irrelevant endurance task.

## Pupil dilation mediated the increased force in the endurance task

In Experiment 2 (N = 30), we aimed at providing physiological evidence that the arousal elevation underlies the facilitation effect due to multitasking observed in Experiment 1. It is well known that performance in both motor and cognitive tasks is modulated by arousal, which is reliably reflected in pupil dilation during the tasks^13,19–24^. Thus, we hypothesized that the concurrent engagement in the endurance and cognitive tasks can elevate an overall arousal level and therefore further increase pupil dilation.

The procedure of Experiment 2 was identical to that of Experiment 1, except that we recorded participants’ pupil size during the experiment (see Methods for details). As with endurance performance, we calculated percent changes in pupil size relative to the PO condition as a baseline due to substantial inter-individual variability in pupil size. As expected, the pupil size changes significantly increased with the difficulty of the cognitive task (Figure 2A; paired t-test, *t*_29_ = 11.210, *p* < 0.001). Results of subsequent tests (one-sample t-test with Bonferroni correction for multiple comparisons) indicated significant increases in pupil size for the 1B (*t*_29_ = 6.686, corrected *p* < 0.001) and 2B (*t*_29_ = 9.723, corrected *p* < 0.001) conditions. These results are consistent with the hypothesis that the demanding cognitive task effectively modulated pupil-linked arousal under the multitask setting.

**Figure 2.**
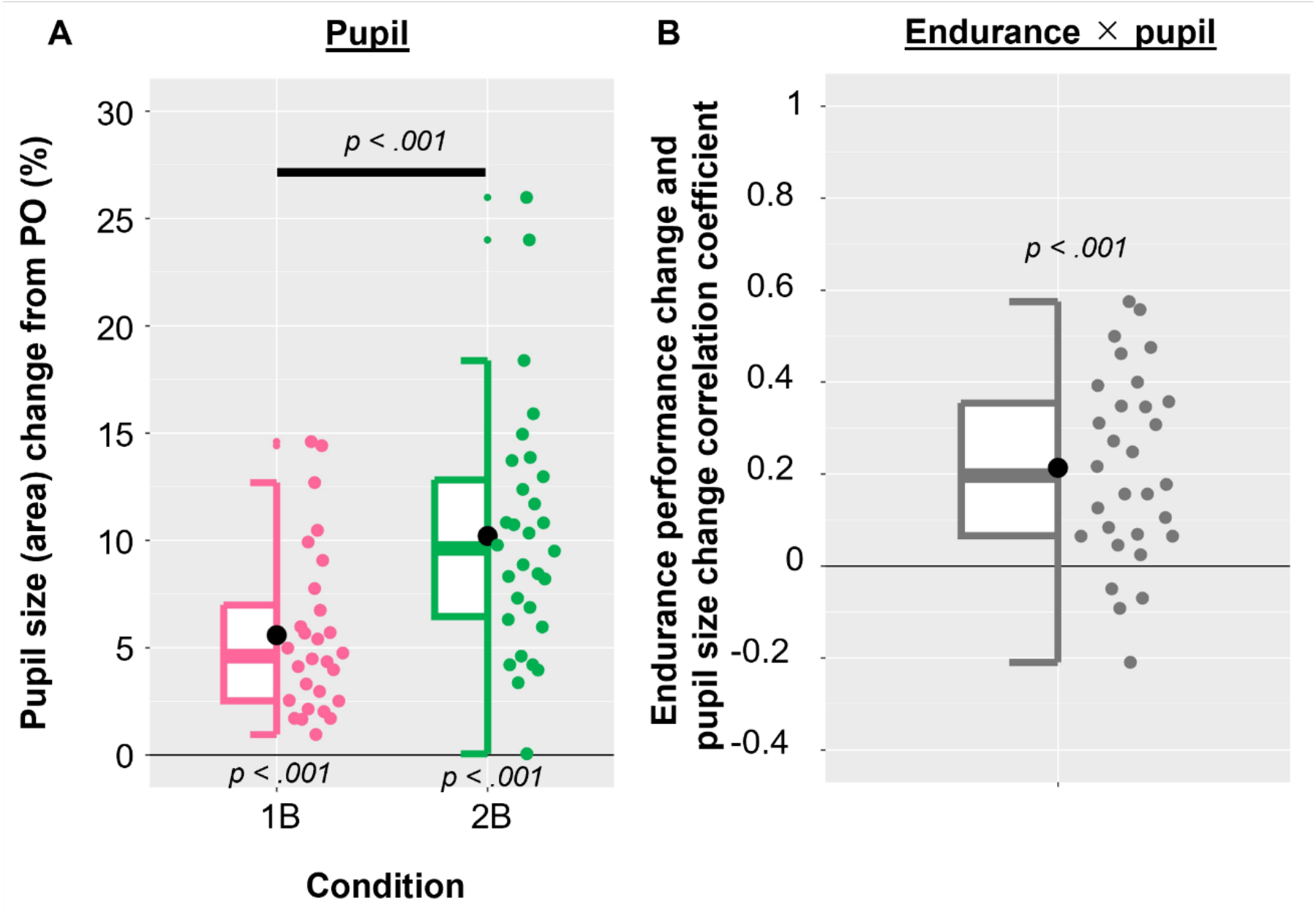
Results of Experiment 2. (A)Changes in pupil size in the 1B and 2B conditions relative to the PO condition as a baseline. Pupil size significantly increased in the 1B and 2B conditions, relative to the PO condition. Pupil dilation was significantly larger in the 2B condition than in the 1B condition. (B)Trial-wise correlation between changes in pupil size and endurance performance. Trial-by-trial fluctuations in pupil size due to the cognitive task correlated with those in the endurance performance within participants. Box plots show the medians, lower and upper quartiles, and minimum and maximum adjacent values without outliers across participants. Black dots represent the mean values across participants, and colored dots represent individual values.

If the increased pupil dilation reflects the elevation of arousal, it should mediate the increased endurance performance. Specifically, we should observe trial-wise correlation between the pupil size changes due to the cognitive task and corresponding changes in the endurance performance. To test this prediction, for each participant we normalized trial-by-trial pupil sizes relative to that in the PO condition as a baseline. Specifically, pupil size on each individual trial in the 1B and 2B conditions was subtracted from the mean pupil size in the PO condition and divided by the standard deviation of pupil size calculated across all trials in the PO condition. Endurance performance on individual trials in the 1B and 2B conditions was normalized in the same way using the mean and standard deviation of endurance performance in the PO condition.

Changes in pupil size and endurance performance were positively correlated for most participants (Figure 2B). Correlation coefficients ranged from -0.209 to 0.575 across participants, and the mean correlation coefficient (0.214) was significantly higher than zero (one-sample t-test; *t*_29_ = 5.755, *p* < 0.001). These results support the hypothesis that the pupil-linked arousal underlies the facilitation effect of multitasking on the endurance performance.

### Reward for the cognitive task also facilitated endurance performance

In Experiment 3 (N = 17), we further tested the validity of the arousal account. In Experiment 2, we manipulated the difficulty of the cognitive task to change pupil dilation (Figure 2). Arousal is a general process modulated not only by task difficulty but also by other factors such as task reward^39–42^. Thus, if the facilitation effect is indeed mediated by arousal, the manipulation of reward for the cognitive task should also lead to a change in pupil dilation and corresponding facilitation effect on endurance performance. Motivation to the cognitive task can also be increased by reward. However, increased motivation does not predict a facilitation effect on endurance performance as it is considered to be a goal-directed processing^31–34^.

Specifically, Experiment 3 used monetary reward to manipulate participants’ task engagement, following previous studies^39–43^. The procedure of Experiment 3 was identical to that of Experiment 2 with the following exception (Figure 3A). Whereas the same PO condition was employed as in Experiments 1 and 2, the other conditions were substituted with two 2-back task conditions with low monetary reward (LR) and high monetary reward (HR). As the task difficulty was identical between the LR and HR conditions, any difference in performance of the endurance task between them should be attributed to the difference in the amounts of reward.

**Figure 3.**
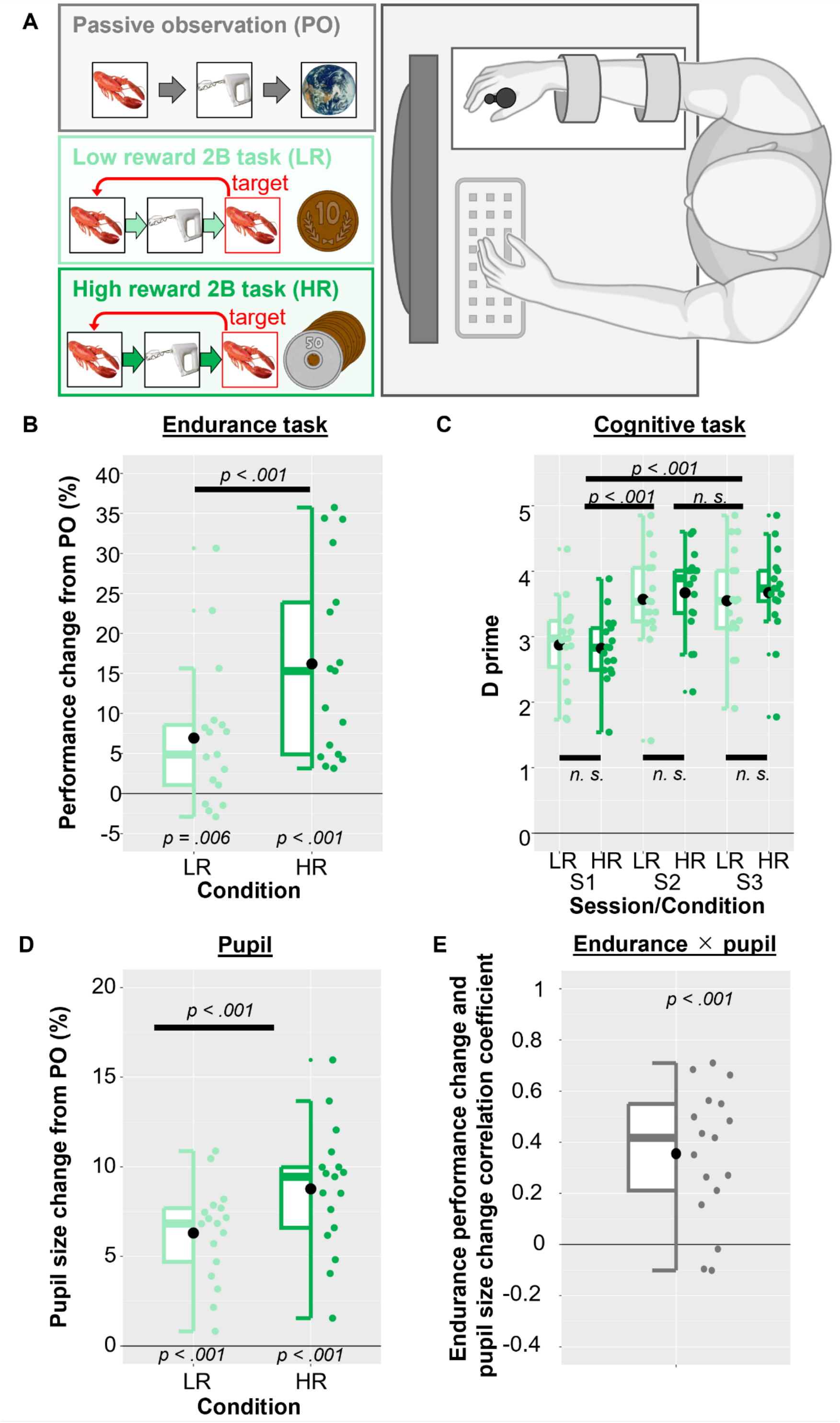
Procedure and results of Experiment 3. (A) Multitask setting in Experiment 3. Participants performed a handgrip muscular endurance task (right) while simultaneously engaging in cognitive task under one of the three conditions (left): passively observing images (PO), performing 2-back task with low reward (LR), and performing 2-back task with high reward (HR). (B) Percent changes in endurance performance in the LR and HR conditions, relative to the PO condition as a baseline. Average endurance performance change relative to the PO condition was significantly higher in the HR than the LR conditions. (C) Cognitive task performance in the sessions S1, S2, and S3. (D) Average pupil size change relative to the PO condition was significantly higher in the LR than the HR conditions. (E) Trial-by-trial fluctuations in pupil size in the LR and HR conditions correlated with trial-by-trial fluctuations in endurance performance within participants. Box plots show the medians, lower and upper quartiles, and minimum and maximum adjacent values without outliers across participants. Black dots represent the mean values across participants, and colored dots represent individual values.

We found that the manipulation of the reward amounts led to increase in the endurance performance, as in Experiments 1 and 2. First, endurance performance change in the LR (one-sample t-test with Bonferroni correction for multiple comparisons; *t*_16_ = 3.161, corrected *p* = 0.012) and HR (*t*_16_ = 5.577, corrected *p* < 0.001) conditions was significantly higher than that in the PO condition (Figure 3B). Second, performance in the HR condition was significantly higher than that in the LR condition (paired t-test, *t*_16_ = 5.845, *p* < 0.001). Third, two-way repeated-measures ANOVA on cognitive performance with the factors of Condition (LR vs. HR) and Session (first multitasking, second multitasking, vs. single task sessions) revealed a significant main effect of Session (*F*_2,32_ = 24.279, *p* < 0.001; Figure 3C). However, no significant main effect of Condition (*F*_2,32_ = 0.711, *p* = 0.412) or interaction between the factors (*F*_2,32_ = 0.493, *p* = 0.615) was found. Thus, levels of cognitive performance matched between the LR and HR condition. Fourth, pupil size changes significantly increased with the amount of reward given to the cognitive task (Figure 3D; paired t-test, *t*_16_ = 4.995, *p* < 0.001). Pupil size changes in the LR (one-sample t-test with Bonferroni correction for multiple comparisons; *t*_16_ = 9.694, corrected *p* < 0.001) and HR (*t*_16_ = 10.256, corrected *p* < 0.001) conditions was significantly larger than that in the PO condition. Thus, additional pupil dilation occurred as the amount of the reward for the 2-back task increased. Finally, we tested whether the increase in endurance performance was mediated by additional pupil dilation using the same normalization method as in Experiment 2. Trial-by-trial changes in pupil size and endurance performance in the LR and HR conditions were significantly correlated within individual participants (range of correlation coefficients: -0.102–0.710; Figure 3E). The mean correlation coefficient (0.355) across participants was significantly higher than zero (one-sample t-test; *t*_16_ = 5.624, *p* < 0.001). These results further support that the facilitation effect of multitasking on the endurance performance occurred due to increase in overall arousal by the concurrent cognitive task beyond the level attainable by the endurance task alone.

## Discussion

The prevailing view on multitasking is that it impairs performance levels in either or both tasks due to interference due to the limited capacity for the tasks^13–16^. The present study challenged this view and revealed the facilitation effect of multitasking on performance. We found that the force exerted in the muscular endurance task increased with the difficulty level and reward for the concurrent cognitive task (Experiments 1 and 3). This facilitation effect was mediated by the additional pupil dilation due to the concurrent tasks (Experiments 2 and 3). The facilitation effect cannot be attributed to focused attention, effort, or motivation to the cognitive task as these processes are task-specific^26–36^. Rather, these results suggest that the concurrent execution of endurance and cognitive tasks elevates overall arousal to a level unattainable by the endurance task, which in turn facilitates endurance performance. To our knowledge, this is the first demonstration that multitasking facilitates task performance.

The present study differed from most research on multitasking in that we used tasks that were previously demonstrated to have little interference^25,37^. Therefore, one might argue that the absence of a deterioration effect in our study would be expected and trivial. However, we deliberately chose these non-interfering tasks to reveal the facilitation effect of multitasking, as interference between tasks has likely masked or canceled out any potential facilitation effect in the previous studies. Importantly, even when a pair of non-interfering tasks are used in previous studies, a common finding is that multitasking had little deterioration effect on performance in each task^25,37^. In contrast, our results show significant facilitation effects on the endurance performance without any cost to the cognitive performance. Our findings are not simply explained by the absence of interference but instead can be attributed to a distinct mechanism by which overall arousal is elevated due to multitasking.

Unlike the facilitation of endurance performance, performance in the cognitive tasks was not facilitated by multitasking (Figure 1C). One possible explanation is the ceiling effect: indeed, d prime values in our experiments were quite high. Another possibility is that arousal levels during multitasking were primarily driven by the difficulty and reward associated with the cognitive task while the endurance task had little impact on arousal. Future studies could examine these possibilities by using a wider range of task difficulties to avoid ceiling effects, and by alternating single-task endurance and cognitive sessions with multitasking sessions to compare arousal levels across different task setups.

While we attributed the facilitation effect to the elevation of arousal, other explanations might be possible. For example, a difficult task or a reward given for a task may increase attentional vigilance or elicit positive emotions. Previous research suggests that such factors influence physical performance^44,45^. However, these factors are known to modulate arousal, as demonstrated by pupillometry studies^46–49^. Thus, even if changes in emotion or vigilance occurred in our study, the common pathway through which they could enhance endurance performance is arousal. At the same time, each of these factors may be related to different forms of arousal, such as emotional and cognitive arousal. It remains debated whether these forms of arousal represent separate processes or different manifestations of the same underlying state^50–52^. Future research may address this issue by disentangling which of these factors predominantly contribute to the elevation of pupil-linked arousal that underlies the facilitation effect of multitasking on performance.

In summary, the present study revealed the facilitation effect of multitasking and its underlying mechanism driven by arousal. Our findings call for a reconsideration of the functional significance of multitasking in daily life and applied settings. Multitasking is not always detrimental to performance; under specific circumstances, two tasks are better than one.

## Methods

### Participants

We recruited healthy adult volunteers (18–54 years old) with normal or corrected-to-normal vision. After exclusions (see Exclusion of participants for details), the number of participants in each experiment was: 24 in Experiment 1 (9 females and 15 males, all right-handed), 30 in Experiment 2 (12 females and 18 males, one left-handed, one ambidextrous), and 17 in Experiment 3 (8 females and 9 males, all right-handed). Handedness was assessed both through self-report and Edinburgh Handedness Inventory test in a post-experiment questionnaire. The experimental protocols were approved by the ethics committee at RIKEN. Written informed consent was obtained from all participants prior to their participation and all participants received monetary compensation for their participation.

### Stimuli and apparatus

Participants were seated in front of an LED monitor with 1,920 × 1,080 resolution and 60 Hz refresh rate. Responses were collected with a keyboard and a force sensor (Current Design) with maximum input of 500 N. During the endurance task, participants positioned their hands on custom-made arm rests. In Experiments 2 and 3, we collected pupil size data with Eyelink (SR Research, Mississauga, Ontario, Canada). In these experiments, a chin rest was used to maintain participants’ head position. Feedback sounds were presented with speakers.

Stimulus presentation was controlled via Psychophysics Toolbox^53,54^ in MATLAB (The MathWorks). Image stimuli extending 4 by 4 degrees of visual angle were presented on a uniform gray background. There were 80 different color images of everyday objects and animals. A green frame with a thickness of 0.5 DVA was presented along the monitor edge as a signal to grip. A 300 ms beep sound was presented as feedback sound.

### Procedure

Each trial started with instructions about the condition, followed by a countdown to the beginning of the task. Participants performed a muscular endurance task in which they were required to grip a force sensor with their hand and sustain the highest force they could for 12 seconds. Simultaneously, they performed one of the three cognitive tasks: passively observing images of everyday objects and animals, 1-back task, or 2-back task, requiring participants to detect images that matched the image presented 1 (1-back task) or 2 (2-back task) images before. Fifteen images were presented on every trial, with four of them being targets in the 1- and 2-back tasks. Each image was presented for 200 ms with 600 ms intervals between the images. Participants had to press a key when they detected a target within 750 ms after its onset. Images were chosen at random on each trial in Experiment 1. In Experiments 2 and 3 images were matched across conditions: 10 sets of randomly chosen images were generated for each session, and the order of images within each set was rearranged to correspond to each of the three tasks. Feedback sounds denoted false alarms and misses in the 1-back and 2-back tasks.

Each session started with three measurements of the participant’s highest force, after which participants performed 30 trials of tasks. These were grouped in mini blocks, each consisting of three trials of different conditions presented in a random order within the mini block. Participants performed two such sessions with order of hand used to grip counterbalanced between them, then performed one baseline session. Baseline session had 10 trials of the 1-back and 2-back task each with the procedure identical to multitasking sessions.

Experiment 3 had different conditions: passive observation, 2-back task with low reward and 2-back task with high reward. Participants were informed that they could earn up to 10 and 90 yen respectively on each 2-back trial depending on their accuracy. They were only informed of the total amount earned at the end of the experiment.

### Exclusion of participants

We excluded participants whose results (percent change of exerted force in the 1B and 2B conditions relative to the PO condition) were outliers based on the following definition: values that are further from median than three median absolute deviations scaled by the inverse normal probability distribution. Consequently, no participants were excluded in Experiment 1, and one participant was excluded in each of Experiments 2 and 3.

### Statistical analysis

To analyze the data from endurance task, we first averaged the exerted force during trial, except for the first second of the trial. We then excluded first three trials of each session and outlier trials, defined as trials on which mean force was more than three standard deviations away from the average force in that condition in that session. To analyze the pupil data, we interpolated the periods of missing data due to blinks and technical issues and excluded the trials on which more than 20% of data had to be interpolated. All the statistical tests performed in this study were two-tailed. The significance threshold was set to an alpha level of 0.05. Analyses of variance (ANOVAs) were performed using the anovakun function in the R statistical software package. For ANOVA, we applied Mendoza’s multi-sample sphericity test to evaluate the sphericity assumption. When necessary, we adjusted the degrees of freedom using Greenhouse-Geisser’s ε to correct any violations.

## Acknowledgements

This work was supported by the RIKEN Special Postdoctoral Researcher (SPDR) program (to S.N. and E.O.), JSPS Kakenhi Grants 19H01041 and 20H05715, and JST Moonshot R&D Grant JPMJMS2013 (to K.S.).

## Author Contributions

All authors conceived and designed the experiments and wrote the manuscript. S.N. conducted the experiments and analyzed data.

## Supplementary information

**Figure S1.**
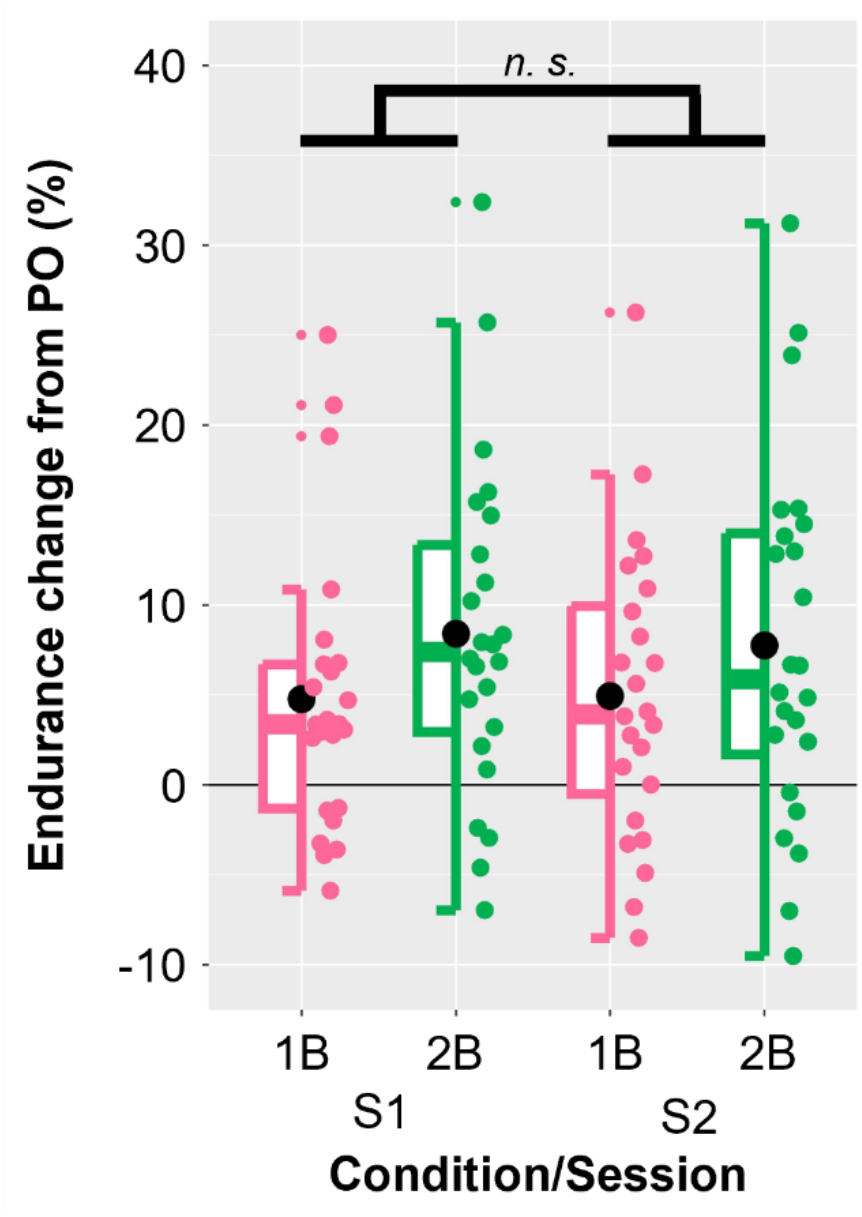
Supplementary results of Experiment 1 with comparison between two multitasking sessions: neither the effect of Session (*F*_1,23_ = 0.014, *p* = 0.907), nor interaction between Session and Condition (*F*_2,46_ = 0.337, *p* = 0.567) were significant, as determined by two-way repeated-measures ANOVA. Box plots show the medians, lower and upper quartiles, and minimum and maximum adjacent values without outliers across participants. Black dots represent the mean values across participants, and colored dots represent individual values.

